# OsIQD14 regulates rice grain shape through modulating the microtubule cytoskeleton

**DOI:** 10.1101/275552

**Authors:** Baojun Yang, Jos R. Wendrich, Bert De Rybel, Dolf Weijers, HongWei Xue

## Abstract

Cortical microtubule (MT) arrays play a critical role in plant cell shape determination by defining the direction of cell expansion^1-3^. The control of plant organ shape and architecture is a major target of cereal crop improvement. Given the pleiotropic effects of MT modification, however, it is challenging to exploiting MT array organization for crop improvement. Moreover, as plants continuously adapt cell growth and expansion to ever-changing environmental conditions, multiple environmental (e.g. light^4^) and developmental (e.g. hormones^5,6^) inputs need to be translated into changes of the MT cytoskeleton. Here, we identify and functionally characterize an auxin-inducible and MT-localized protein OsIQ67-DOMAIN14 (OsIQD14), which is highly expressed in rice seed hull cells. While deficiency of *OsIQD14* results in short and wide seeds and increases overall yield, overexpression leads to narrow and long seeds, caused by changes in the direction of MT arrangement. We further show that OsIQD14-mediated MT reordering is regulated through interacting with SPIRAL2, a MT-binding protein involved in KATANIN1-mediated MT rearrangement^7,8^, and with calmodulin proteins. As such, OsIQD14 acts as an integrator of auxin and calcium inputs into MT rearrangements, and allows effective local cell shape manipulation to improve a key rice yield trait.

---

The IQ67 domain (IQD) protein family contains a conserved 67 amino acid domain, and was first identified in Arabidopsis, where it is represented by a gene family with 33 members^9^. Several IQD proteins have been shown to interact with Calmodulin through their IQ67 domain^10-12^. However, the biological relevance of this interaction and the function of IQD proteins remains largely unknown. The AtIQD15-18 subclade was suggested to act downstream of the AUXIN RESPONSE FACTOR5/MONOPTEROS (ARF5/MP) transcription factor^13^. Rice shares OsIQD14 as a single ortholog to the AtIQD15-18 subclade (Wendrich et al, submitted^14^). To investigate the role of *OsIQD14*, we first analyzed its expression pattern using the transcript digital gene chip and found it to be highly expressed in inflorescences, pistils, and spikelet hull tissues (Supplemental Fig. 1a-b). Quantitative RT-PCR (qRT-PCR) analysis showed that *OsIQD14* transcripts can be detected during panicle development, peaking around the middle stage and gradually decreasing in mature stages of development (Supplemental Fig. 1c). We further analyzed the expression pattern by fusing the *OsIQD14* promoter region to the ß-glucuronidase (GUS) reporter. Young spikelet hulls and anthers showed a strong GUS signal (Fig. 1a-c), indicating that *OsIQD14* may play a role during rice spikelet hull development. Because members of the *AtIQD15-18* subclade act downstream of MP^13^ and are auxin-inducible (Wendrich et al, submitted^14^); we analyzed *OsIQD14* transcript levels upon exogenous auxin treatment. qRT-PCR analysis confirmed that *OsIQD14* transcripts were quickly induced upon auxin (Indole 3-Acetic Acid) treatment (Fig. 1d). Moreover, the rice ortholog of *AtMP, OsARF11*, is transcribed throughout the panicle development similar to *OsIQD14* (Supplemental Fig. 1d), suggesting that the auxin-inducible *OsIQD14* transcripts could be under the control of OsARF11.

**Figure 1:**
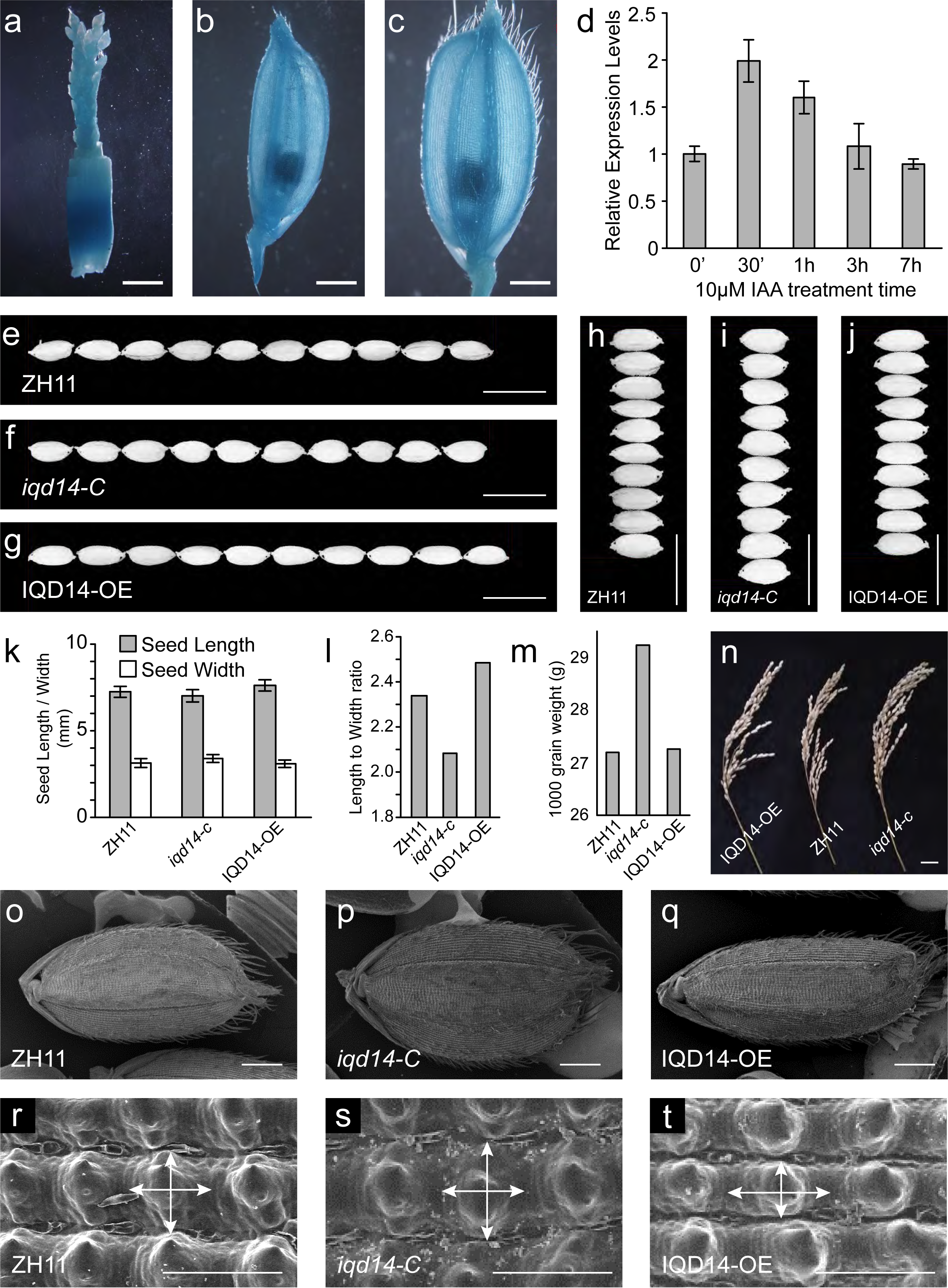
*OsIQD14* regulates rice seed size and hull cell shape. **a-c,** Promoter-GUS fusion studies showed the *OsIQD14* expression in young panicle (0.4 cm) and lemma before anthesis of rice. Representative images are shown. Bar=2mm. **d,** qRT-PCR analysis revealed the up-regulated expression of *OsIQD14* under short time (30min-420 min) auxin treatment (10 μM IAA). Rice seedling roots were used and expressions were normalized with the *ACTIN* transcript and relative expression levels were calculated by setting the *OsIQD14* expression in the absence of auxin as “1.0”. Experiments were biological repeated and data are shown as mean ± standard error (SE). **e-j,** Grain morphology of ZH11 and rice transgenic plants deficiency of (*iqd14-C*) or overexpressing (p35S::*IQD14* or *IQD14-OE*) OsIQD14. Bar=20 mm. **k-m,** Measurement and statistical analysis of seed length and width (**k**), length/width ratio (**l**), and thousand-grain-weight (**m**) of ZH11, *iqd14-C* and *IQD14-OE* plants. data are shown as mean ± SE (n>300). **n,** Harvested panicles of one plant individual in ZH11, *iqd14-C* and *OsIQD14-OE* plants. Scale bar=2 cm. **o-t,** Scanning electron microscopy observation of seeds with hulls and outer glume (**r-t,** bar=100 μm) of the lemma among ZH11, *iqd14-C* and *IQD14-OE* plants. Bar=1 mm.

To define the biological function of *OsIQD14*, we generated loss-of-function mutants by using CRISPR/Cas9 technology^15^. Construct expressing a guide RNA targeting the first exon of *OsIQD14* was transformed into ZH11 wild type rice plants. Four independent homozygous lines were isolated that carried frame shift mutations resulting from a 1-bp insertion or a 5-bp, 22-bp or 34-bp deletion, respectively (Supplemental Fig. 2a). None of the independent mutants presented a visible phenotype during vegetative growth, but all produced wider and shorter grains compared to those of ZH11 (Fig. 1e-f, h-i and Supplemental Fig. 2b), demonstrating a role for *OsIQD14* in panicle and spikelet development. For all further analyses, the 5-bp deletion *iqd14-c* mutant was selected. Next, we generated overexpression plants by driving *OsIQD14* from the strong p35S promoter in a ZH11 background (p35S::*OsIQD14* or *OsIQD14-OE*). In contrast to the ZH11 and *iqd14-c* mutant, *OsIQD14-OE* plants produced narrower and longer grains (Fig. 1g, j and k-l). Furthermore, the 1000-grain weight of *iqd14-c* was significantly increased compared to that of ZH11, while that of *OsIQD14-OE* was similar to ZH11 (Fig. 1m). Importantly, the panicle of *iqd14-c* and *IQD14-OE* was similar to that of ZH11 (Fig. 1n). As the spikelet hull has been proposed to restrict growth of a grain and as such determines the grain size of rice, we examined cell morphology in the outer glume of the lemma of ZH11, *iqd14-c* and *IQD14-OE* plants. Scanning electron microscopy (SEM) analysis revealed that *iqd14-c* spikelet hull cells were shorter and wider than that of ZH11. Conversely, the hull cells of *IQD14-OE* plants were narrower and longer than those of ZH11 and *iqd14-c* plants (Fig. 1o-t and Supplemental Fig. 2c-e and 3a-i). Taken together, these results indicate that the changes in overall seed shape by modifying *OsIQD14* levels (e.g. short and wide versus long and slender) are caused by the same type of changes to the individual spikelet hull cells, and that modifying *OsIQD14* transcript levels can be used as a tool to modify rice grain shape without a yield penalty under normal growth conditions.

**Figure 2:**
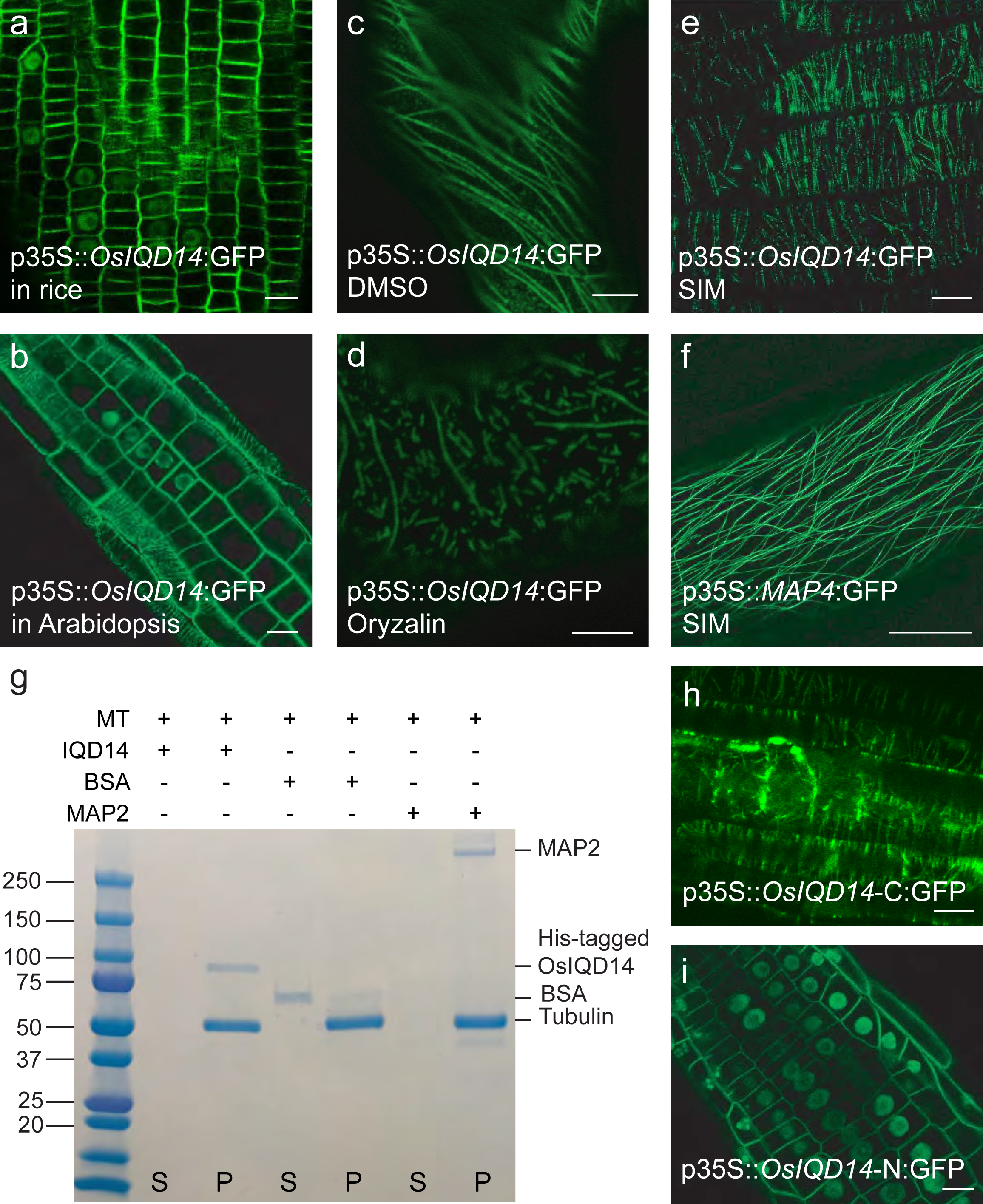
OsIQD14 protein is located at microtubules and in the nucleus **a,** Subcellular localization of 35S::*OsIQD14*:GFP in rice root meristem region. Bar=10μm. **b,** Subcellular localization of 35S::*OsIQD14*:GFP in *Arabidopsis* root meristem region (Bar = 10μm). **c-d,** Super resolution structured illumination microscopy (SIM) analysis of 35S::*OsIQD14*:GFP and p35S::*MAP4*:GFP localization at microtubules in *Arabidopsis thaliana* root epidermis cells at meristem region (Bar = 5μm) (**c**) and 10 μm (**d**). **e-f,** IQD14 protein is localized to microtubules (**e**) and is depolymerized by 50μM oryzalin treatment for 1.5 hour (**f**) in *Nicotiana benthamiana* epidermis cells (Bar = 5μm). **g.** *In vitro* microtubule spin down assay showed that OsIQD14 binds to microtubules directly. IQD14-His protein was incubated without or with taxol-stabilized bovine MT (20 µM), divided into pellet and soluble (Supernatant) fractions by ultracentrifugation, and then separated by 4-12% SDS-PAGE and stained with Coomassie brilliant blue. MAP2: positive control protein (280 kD), which bind to microtubules. BSA: negative control protein. 68 kD. **h-i,** Root cells of 7 days old transgenic *Arabidopsis* seedlings expression C terminal of IQD14-GFP (**h,** Bar = 5μm) or N terminal of IQD14-GFP (**i,** Bar = 20μm), which localized to microtubules or nucleus, respectively.

To understand through what mechanism OsIQD14 can alter the cell shape, we first analyzed its subcellular protein localization. Similar to some IQD protein in *Arabidopsis*^10, 11^, p35S::*OsIQD14*:GFP was localized to the nucleus and cytoskeleton-related structures in rice root mersitems (Fig. 2a), *Nicotiana benthamiana* epidermal cells (Supplemental Fig.4a) and *Arabidopsis thaliana* root meristems (Fig. 2b). Considering that the cytoskeleton-associated localization was abolished by application of the MT depolymerization drug oryzalin (Fig. 2c-d), OsIQD14 thus likely localizes to MT. To better understand this MT-associated localization, we used super resolution Structured Illumination Microscopy (SIM) in Arabidopsis root meristem cells. Intriguingly, we observed punctate localization of OsIQD14 along MT (Fig.2e), while structural MT proteins (p35S::GFP:*TUA6*^16^) or general MT associated proteins (p35S::*MAP4*:GFP^16^) showed uninterrupted distribution along the filaments (Fig. 2f). This result suggests that OsIQD14 is not a generic MT binding protein, but could potentially regulate MT behavior at specific positions. Although nuclear localization of IQD proteins has been reported previously^10^, OsIQD14 showed strongly dynamic nuclear localization that depends on the cell cycle state (Fig. 2b and Wendrich et al, submitted^14^), suggesting that IQD function during cell division is complex.

To examine whether OsIQD14 interacts with MT directly, we performed a MT spin down assay. Similar to MAP2, a generic MT-binding protein and positive control^17^, OsIQD14 was detected in the pellet fraction (Fig. 2g), demonstrating that OsIQD14 directly binds MT. To elucidate which part of OsIQD14 localized to MT, we generated GFP fusion proteins consisting of only the N- or C-terminal region of OsIQD14. Although both MT and nuclear localization was observed in the p35S::*OsIQD14*:GFP line expressing the full-length protein (Fig. 2b), the C-terminal region of OsIQD14 fused to GFP showed MT localization only, while the N-terminal region of OsIQD14 fused to GFP showed nuclear localization only (Fig. 2h-i and Supplemental Fig. 4d-i). Taken together, these results show that IQD14 binds MT directly through its C-terminal domain.

Based on the combined results described above, it is tempting to speculate that OsIQD14 controls spikelet hull cell shape by modifying the MT cytoskeleton. Unfortunately, the hardened spikelet hull cells in rice are unsuited for observing MT ordering and dynamics. However, given the identical protein localization and evolutionary conservation between AtIQD18 and OsIQD14 (Wendrich et al, submitted^14^), we transformed p35S::*OsIQD14*(*OsIQD14*-OE) in *Arabidopsis*. Similar to *AtIQD16*^10^ and *AtIQD18*(Wendrich et al, submitted^14^) overexpression, *Arabidopsis OsIQD14-OE* seedlings showed narrow, long and spiraling cotyledons (Fig. 3a-b). Also, epidermal pavement cells became largely isodiametric and lost the typical jigsaw puzzle shape found in Col-0 control plants (Fig. 3c-d and Supplemental Fig. 2f-g). Indeed, MT topology was strongly affected in *OsIQD14-OE* plants compared to control plants as visualized by using p35S::*MAP4*:GFP (Fig. 3e-f). In conclusion, these results indicate that *OsIQD14* overexpression in *Arabidopsis* alters the MT orientation. By inference, this suggests that OsIQD14 controls cell shape through affecting the MT cytoskeleton.

**Figure 3:**
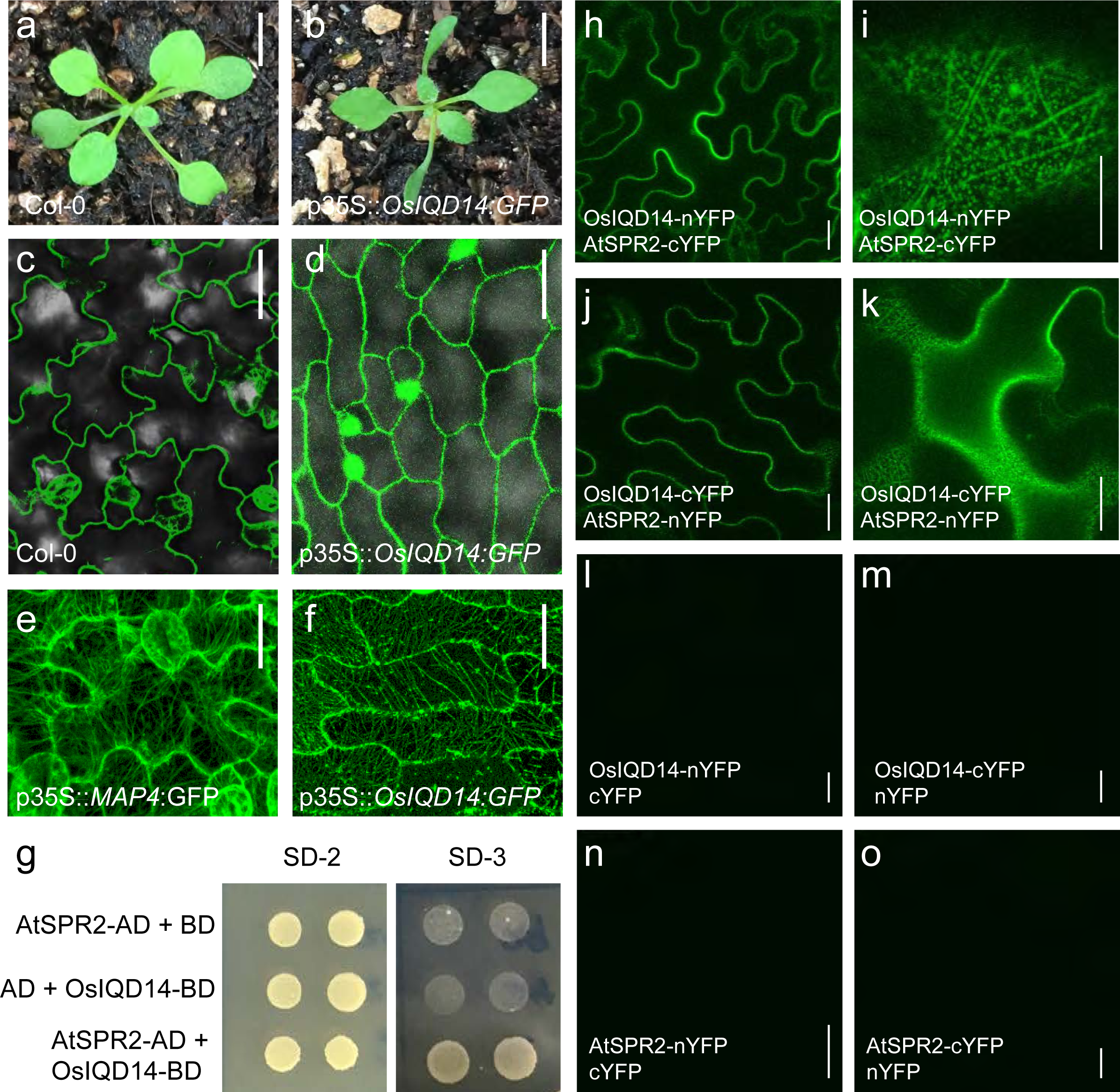
OsIQD14 regulates the shape of *Arabidopsis* pavement cells by affecting microtubule ordering. **a-b,** Phenotype of wild type (Col-0) and p35S::*OsIQD14* in *Arabidopsis*(Bar = 0.5cm). **c-d,** Epidermal pavement cell phenotype of Col-0 and p35S::*OsIQD14*(Bar = 25μm). **e-f,** Microtubule ordering in p35S::*MAP4*:GFP and 35S::*OsIQD14*:GFP pavement cells (Bar = 20 μm). **g,** Yeast-2-hybrid assay showed that OsIQD14 interacts with SPR2 directly *in vitro*. Full-length cDNAs of OsIQD14, SPR2 were subcloned into pGBKT7 and pGADT7 vectors and yeast cells were grown on synthetic dropout (SD-2: -Trp, -Leu; SD-3: -Trp, -His, -Leu) medium. Empty pGADT7 or pGBKT7 vectors transformed with OsIQD14-BD and SPR2-AD were used as negative controls. AD, activating domain; BD, binding domain. **h-o,** BiFC assay showed that the OsIQD14 interact with SPR2 at microtubule in *Nicotiana benthamiana* epidermis cells by co-expression of IQD14-n/cYFP and SPR2-c/nYFP (Bar = 20μm). An enlarged view was showed as **i** and **k** (Bar = 10μm). Empty n/c YPF was used as negative controls (**i-o**).

Intriguingly, the phenotypes observed upon *OsIQD14* expression in *Arabidopsis* seedlings are very similar to those seen in the *spiral2* mutant^8,18^. SPIRAL2 (SPR2) is a plant-specific MT binding protein required for anisotropic growth in *Arabidopsis*^8,18-20^. It was shown to protect MT minus ends to promote KATANIN-dependent severing and reorientation of plant cortical microtubule arrays^7^. Given this striking phenotypic resemblance, we focused our attention on SPR2 in order to further understand how OsIQD14 could affect MT organization. We first tested a possible interaction between OsIQD14 and AtSPR2 using yeast two hybrid (Y2H) and bimolecular fluorescence complementation (BiFC) analyses (Fig. 3g-o). Both the *in vitro* Y2H and the *in vivo* BiFC showed that OsIQD14 could interact with AtSPR2. Moreover, an interaction between the orthologous AtIQD18 and SPR2 was also recently identified (Wendrich et al, submitted^14^), strengthening these findings and suggesting that IQD-SPR2 interactions are a conserved property. Interestingly, the OsIQD14-SPR2 interaction using BiFC localized to punctuated regions on MT-like structures (Fig. 3i); suggesting IQD14 could affect MT dynamics by binding to and regulating SPR2 activity.

Although binding of Calmodulin to the IQ67 domain of IQD proteins has been reported^10-12^, the biological significance of this binding remains unclear. We thus tested the interaction between OsIQD14 and three rice Calmodulins, named OsCaM1, 2 and 3, all of which showed similar expression patterns to *OsIQD14* in digital gene expression analysis (Supplemental Fig. 5a-c). Y2H and BiFC analyses showed that OsCaM1, 2 and 3 interact with OsIQD14 both *in vitro*(Fig. 4a) and *in vivo* at MT structures in tobacco leaf epidermal cells (Fig. 4b-g). Similar to other IQDs in *Arabidopsis*, IQD14 interacting with CaM1 was enhanced by calcium treatment (Fig. 4h). We next aimed to investigate the biological significance of the OsCaM1-OsIQD14 interaction. Introducing a p35S::*OsCaM1* construct into the *Arabidopsis* p35S::*OsIQD14*(*OsIQD14*-OE) plants reverted all phenotypes related to *OsIQD14* overexpression back to Col-0 controls (Fig. 4i-n). These included the long, narrow and spiraling cotyledons, and the absence of jigsaw-shaped epidermal pavement cells (Fig. 4o-r). Taken together, these results strongly suggest that the Ca^2+^-dependent Calmodulin-Os*IQD14* interaction inhibits IQD14 activity. Interestingly, the CaM-binding and MT-localization properties are separable: the CaM-interacting IQ67 domain is located in the N-terminus of OsIQD14, while the C-terminus is sufficient for MT localization (Fig. 2h). Thus, the OsIQD14 protein likely recruits CaM to MT filaments through these two binding modules.

**Figure 4:**
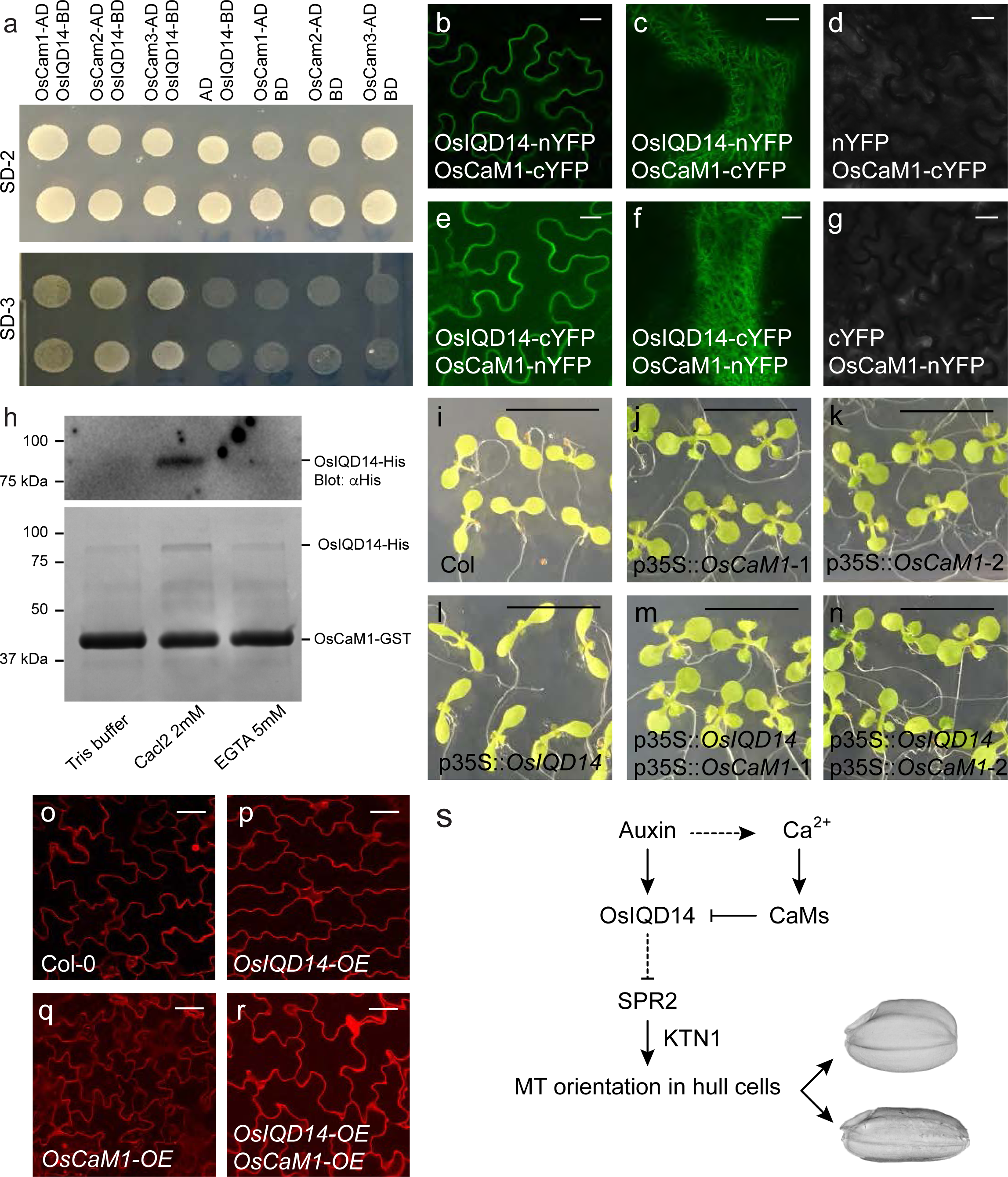
Calmodulins inhibit OsIQD14 activity. **a,** Yeast-2-hybrid assay showed that IQD14 interacts with OsCaM1, 2 and 3 directly *in vitro*. Full-length cDNAs of OsIQD14, OsCaM1/2/3 were subcloned into pGBKT7 or pGADT7 vectors and yeast cells were grown on synthetic dropout (SD-2: -Trp, -Leu; SD-3: -Trp, -His, -Leu) medium. Empty pGADT7 or pGBKT7 vectors transformed with OsIQD14-BD and OsCaM1/2/3-AD were used as negative controls. AD, activating domain; BD, binding domain. **b-g,** BiFC assay showed that OsIQD14 interacts with OsCaM1 at microtubules in *Nicotiana benthamiana* epidermis cells by co-expression of IQD14-n/cYFP and CaM1-c/nYFP,(Bar = 20μm). **c** and **f** showed an enlarged view of interaction zone closed to plasma membrane (Bar = 5μm). Empty n/c YPF was used as negative controls (**d** and **g**). **h,** Calcium dependent OsCaM1-OsIQD14 *in vitro*. GST beads loaded with GST-tagged Calmodulin (CaM1) were incubated with IQD14-His protein at 4°C in the presence of 2 mM CaCl2 or 5 mM EGTA. Pellet (GST-OsCaM1 beads) fraction was detected by probing with His antibody (top) or stained with Coomassie brilliant blue (bottom). **i-n,** Cotyledon phenotype of the indicated genotypes (Bar = 1cm). **o-r,** Epidermal pavement cell phenotype of the indicated genotypes. PI was used to stain cell wall (Bar = 50μm). **f,** A proposed model for Calmodulin-IQD14-SPR2 in regulating microtubule ordering in rice hull cells and seed size.

Being the primary agronomic value trait, grain size and shape are key parameters for rice improvement. A range of regulators of grain size or shape have been identified through studying natural variation or mutagenesis^21, 22^. Identification of causal genes has revealed the role of transcription factors (such as OsSPL16^23^, OsSPL13^24^, OsGRF4^25^), a TRM-containing protein (GW7^23^), an auxin-metabolism protein (TGW6^26^), etc^21^. While these functions are diverse, no coherent mechanistic framework has emerged for the cellular basis underlying the altered morphology. Here, we identify the IQD14 gene as a key regulator of grain shape, acting on cell shape determination through controlling the MT cytoskeleton (Fig. 4s). Intriguingly, while mutations and overexpression consistently affect grain properties, no adverse pleiotropic effects are observed. Thus, this gene allows direct modulation of MT behavior and cell shape in husk cells to affect grain properties without the pleiotropic effects that manipulation of the MT cytoskeleton often induces^27^. Besides the possibility for improving a rice agronomic trait through manipulating the MT cytoskeleton without fitness penalties, we also present a mechanism that explains how multiple external and internal signals, in this case Calcium and auxin, can be integrated by OsIQD14 and translated into modifying MT ordering via SPR2. Given that similar activities and regulation were found for the *Arabidopsis* orthologs of OsIQD14, this module is likely deeply conserved in flowering plants. Interestingly, while molecular mechanisms are not yet resolved, altered expression of a tomato IQD gene (SUN) leads to altered fruit shape^28^. Thus, our study not only reveals a new mechanism for signal integration at the MT cytoskeleton, but also identifies a new role for Ca^2+^ in recruiting CaM proteins to MT and modulating the OsIQD14-SPR2 interaction. As such, our results offer novel mechanisms for organ shape modification in a wide range of crop plants.

## Methods

### Plant materials and growth conditions

Rice (*Oryza sativa*, japonica variety Zhonghua11, ZH11) plants were cultivated in the field at Shanghai under natural growing conditions. For growth of transgenic plants, rice seeds were germinated in sterilized water and grown in a phytotron under a 12 h light (28°C) / 12 h dark (22°C) cycle. *Arabidopsis thaliana* ecotype Columbia-0 (Col-0) was used in all transformation and phenotype analysis. All seeds were germinated on MS (Murashige and Skoog, Duchefa) medium after three days at 4^°^C. Seedlings and plants were grown in a phytotron at 22^°^C with a 16-h light/8-h dark photoperiod.

### Vector construction and plant transformation

The vector CRISPR/Cas9-*iqd14* was generated by using two 20-bp fragments from the first exon of OsIQD14 introducing into pOsCas9 vector, and then this plasmid was transformed into rice (ZH11) by *Agrobacterium tumefaciens*-mediated transformation^29^. The IQD14 cDNA was amplified by PCR with primers IQD14-1 and IQD14-2 (Table S1) using total cDNA of ZH11 seedlings as template and subcloned into pENTR/D-TOPO (Invitrogen) to generate the pENTR/D-TOPO-IQD14 construct. For stable transformation, p35S::*OsIQD14*:GFP was generated by LR reactions with pGWB5 using pENTR/D-TOPO-IQD14, which was then transformed into ZH11. Expression levels of p35S::*OsIQD14*:GFP plants were examined by qRT-PCR and confirmed positive lines were used for further analysis.

Transformation of *Arabidopsis* Col-0 plants was performed by the floral-dipping procedure. Plants expressing p35S::CAM1:RFP and p35S::IQD14:GFP were generated by introducing a p35S::OsCaM1:RFP construct into p35S::OsIQD14 plants. Primers are listed in Supplementary Table S1.

### Promoter-reporter gene fusion studies and GUS activity analysis

To analyze the expression pattern of *IQD14* gene, a 700-bp DNA fragment of *OsIQD14* promoter was amplified by PCR using ZH11 genomic DNA as template and subcloned into pENTR/D-TOPO vector. The resultant construct p*OsIQD14*::GUS was generated by LR reaction with pGWB4 and transformed into ZH11 and confirmed positive lines were used for further analysis. GUS activity of T2 homozygous progeny of independent lines were detected according to previous description^30^ and photographed using a Nikon SMZ 800 stereoscope with a Nikon digital Coolpix 995 camera.

### RNA extraction and quantitative real-time PCR (qRT-PCR) analysis

Total RNAs were extracted using Trizol reagent (Invitrogen) and reversely transcribed to first-strand cDNA. qRT-PCR analysis was performed with Real-Time PCR Master Mix (Toyobo) and data were collected using the Bio-Rad Real Time detection system in accordance with the manufacturer’s instruction manual. Primers were listed in Supplementary Table S1. Expression of *IQD14* was analyzed using primers IQD14-RT1 and IQD14-RT2.

### Scanning electron microscopy observation of spikelet hull

Cell number and cell area in the outer parenchyma layer of the spikelet hulls were measured by Olympus stream software. The sample pretreatment for scanning electron microscopy observation (S-3000N; Hitachi) was performed as described previously^31^. Grain weight were analyzed as described previously^32^.

### Subcellular localization analysis

Fluorescence of transgene seedlings and tobacco epidermal cells was observed by confocal laser scanning microscopy (Leica SP8) with an argon laser excitation wavelength of 488 nm or 561 nm. For imaging with SIM, the Alpha Plan Apochromat 1003, NA 1.57 oil objective was used, and images were acquired from a single optical section.

### Protein-protein interaction assays

Interaction of IQD14 and SPR2 (or CaM1) was detected by standard Y2H analysis following the manufacturer’s instructions (Clontech). cDNAs encoding OsIQD14, AtSPR2, OsCaM1,2,3 were subcloned into pGBKT7 and pGADT7 vector, resulting in the fusion of IQD14-AD, SPR2-AD, CaM1-AD, IQD14-BD, SPR2-BD and CaM1-BD respectively (AD, activating domain; BD, binding domain). Primers are listed in Supplementary Table S1. Yeast transformants were spotted on the restricted SD medium (SD-Leu/-Trp, short as SD-L/T) and selective medium (SD-Leu/-Trp/-His/, short as SD-L/T/H). For BiFC (Bimolecular Fluorescence Complementation) assay, cDNAs encoding *IQD14, SPR2* or *CaM1* were cloned into p35S::YFP-N or p35S::YFP-C vector by gateway LR reaction, resulting in constructs expressing IQD14-cYFP, SPR2-nYFP and CaM1-nYFP, respectively. Resultant constructs with control vectors were co-expressed in *N. benthamiana* leaves and YFP was observed by Leica SP8 confocal microscope using an argon laser excitation wavelength of 488 nm after infiltration for 3 days.

### Recombinant expression of OsIQD14 and MT spin down assay

Coding sequences of *OsIQD14* were amplified by PCR (primers *IQD14-COLD-P1/2*) and subcloned into pCold-HF for expression of OsIQD14-His fusion proteins. After confirmation by sequencing, the construct was transformed into *E. coli* BL21(DE3) cells, and expression of the fusion protein was induced by adding isopropyl-β-D-thiogalactoside (final concentration 1 mM) at 16oC overnight. The cells were lysed by sonication in lysis buffer (50 mM NaH2PO4, 300 mM NaCl and 10 mM imidazole, pH 8.0) and OsIQD14-His protein was purified using Ni-NTA His Bind Resin (Novagen) according to the manufacturer’s protocols. *In vitro* MT binding assay was performed using MT binding protein spin-down assay kit (Cytoskeleton). Briefly, 5 μg purified His-OsIQD14-His protein was incubated with 10 μg prepolymerized bovine brain tubulin in general tubulin buffer (80 mM PIPES, pH 7.0, 2 mM MgCl2, and 0.5 mM EGTA) containing 20 μM taxol followed by centrifugation at 100,000g, and both soluble and pellet fractions were analyzed by SDS-PAGE and Coomassie Brilliant Blue staining.

### Expression of recombinant OsCaM1 and calmodulin binding assay

Calmodulin binding assay were performed according to previous description with some modifications^12^. For expression of CaM1-GST, a full-length cDNA fragment encoding the OsCaM1 were first amplified and then subcloned into pENTR/D-TOPO (Invitrogen). To express the OsCaM1-GST fusion protein, pDEST-GST were used with Gateway LR Clonase II enzyme mix (Invitrogen). The recombinant OsCaM1 protein was expressed in BL21(DE3) at 30^°^C for 4 h by induction with 1 mM IPTG. Bacterial cells were harvested and sonicated in Lysis buffer (50 mM Tris-HCL, 150 mM NaCl). After centrifugation, the supernatant was used for incubating with GST agarose. Aliquots of 100 μL of CaM1-GST beads, pre-equilibrated with Lysis buffer, were mixed with 500 μL of bacterial supernatant supplemented with 2 mM CaCl2 or 5 mM EGTA and incubated for 1 hour at 4 °C under gentle shaking. CaM1 beads were sedimented by centrifugation and washed four times with 500 μL of Lysis buffer, followed by a final wash with 100 μL of the same solution. The bound proteins were eluted by boiling the beads for 2 min in 100μL of 4x SDS sample buffer. Proteins of the total extract, the initial supernatant, the last wash, and the pellet fraction were analyzed by SDS-PAGE and western blot using antibody against His.

## Acknowledgements

Funding for this research is gratefully acknowledged from the National Transformation Science and Technology Program (2016ZX08001006-009), The National Key Research and Development Program of China (2016YFD0100501, 2016YFD0100902) and the National Natural Science Foundation of China (31671660). B.D.R. was funded by The Research Foundation - Flanders (FWO; Odysseus II G0D0515N and 12D1815N) and Netherlands Organization for Scientific Research (NWO; VIDI 864.13.00). We thank Ms. Shu-Ping Xu (SIPPE) for assisting with the rice transformation.

## Author contributions

H.X. and D.W. conceived the project; B.Y. and J.R.W. designed experiments; B.Y. performed experiments; H.X. and D.W. supervised the project; B.Y., B.D.R., D.W. and H.X. wrote the paper with input from all authors.

## Competing interests

The authors declare no competing financial interests.

---

Supplemental Figure legend

**Supplementary Figure 1.**
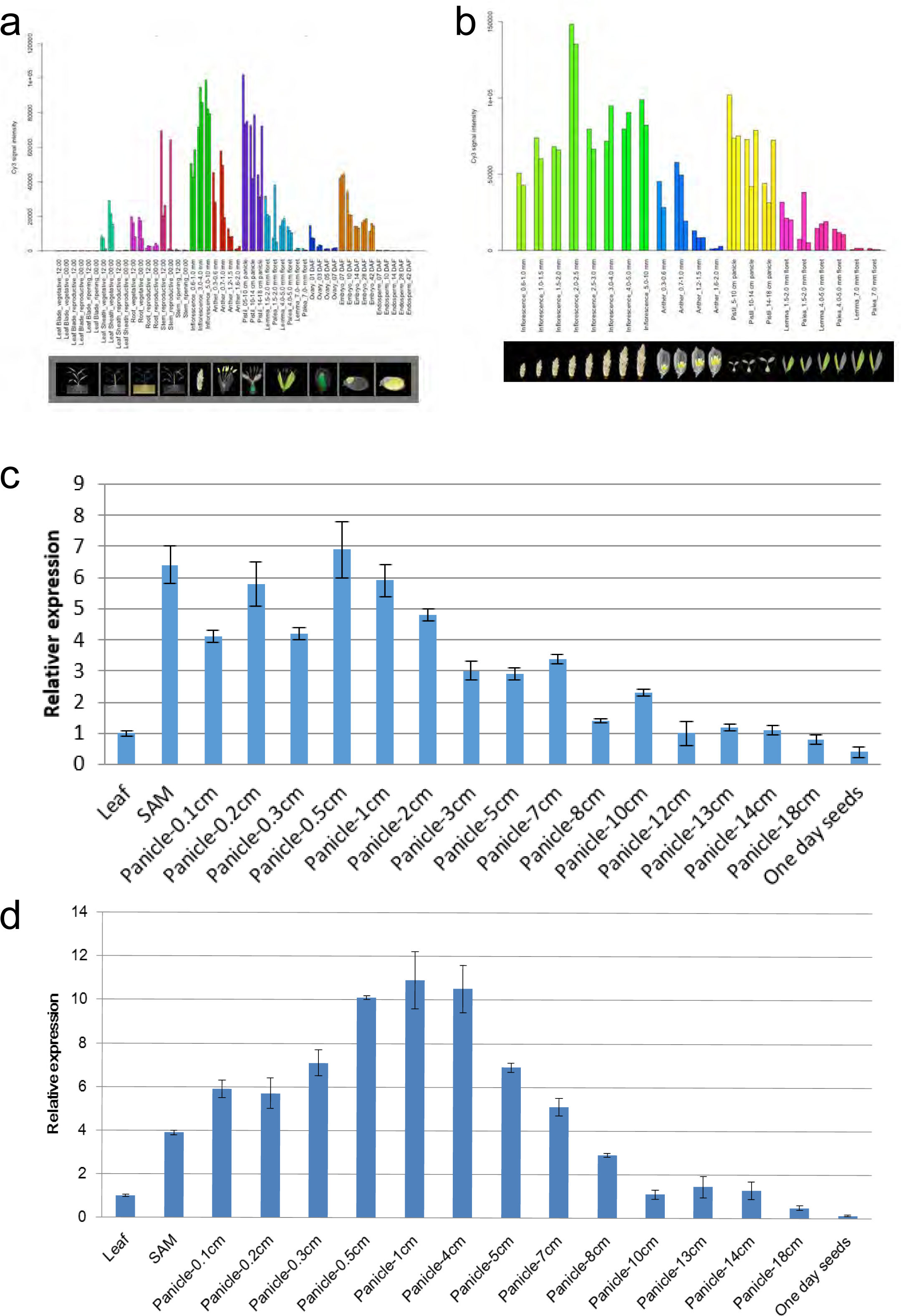
Expression pattern of *OsIQD14* and *OsARF11* in rice tissues. **a,** *OsIQD14* digital expression levels in leaf, root, inflorescence, anther, pistil, lemma, palea, embryo and endosperm. **b,** *OsIQD14* digital expression levels in different stage of inflorescence, anther, pistil, lemma and palea. **c,** Quantitative RT-PCR (qRT-PCR) analysis revealed the transcription of *OsIQD14* in various tissues including SAM, leaf, and young panicles (YP) at different developmental stages (indicated as the lengths of panicles, cm). The expressions were normalized with the *ACTIN* transcript and relative expression levels were calculated by setting the *OsIQD14* expression in Leaf as “1.0”. Experiments were biological repeated and data are shown as mean ± standard error (SE). **d,** Quantitative RT-PCR (qRT-PCR) analysis revealed the transcription of *OsARF11(MP)* in various tissues including SAM, leaf, and young panicles (YP) at different developmental stages (indicated as the lengths of panicles, cm). The expressions were normalized with the *ACTIN* transcript and relative expression levels were calculated by setting the *OsARF11(MP)* expression in Leaf as “1.0”. Experiments were biological repeated and data are shown as mean ± standard error (SE).

**Supplementary Figure 2.**
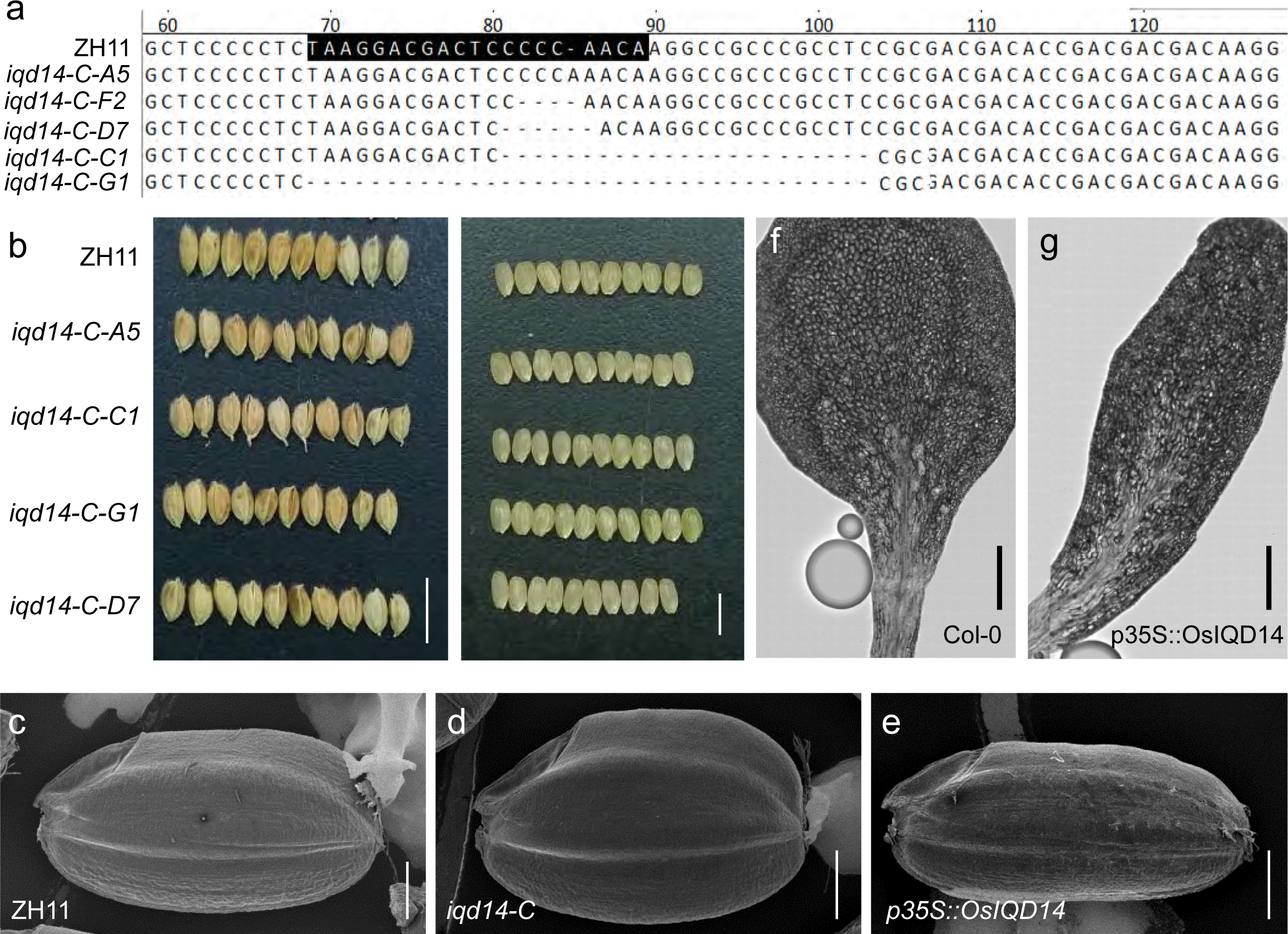
Additional OsIQD14 loss- and gain-of-function phenotypes **a,** Targeted mutagenesis of *IQD14* gene. The position of target site is shown on the gene sequence. The spacer sequence is shaded in black. Sequencing results of mutant alleles are aligned to the reference genome sequence. **b,** Seed phenotype of different mutagenesis of *IQD14*(Bar = 1cm). **c-e,** Scanning electron microscopy observation of seeds without hulls among ZH11 (**c**), *iqd14-C*(**d**)and p35S::*OsIQD14*(**e**) plants (Bar = 1mm). **f-g,** Cotyledon phenotype of Col-0 (**f**) and p35S::*OsIQD14* plants (**g**) (Bar = 1.5mm).

**Supplementary Figure 3.**
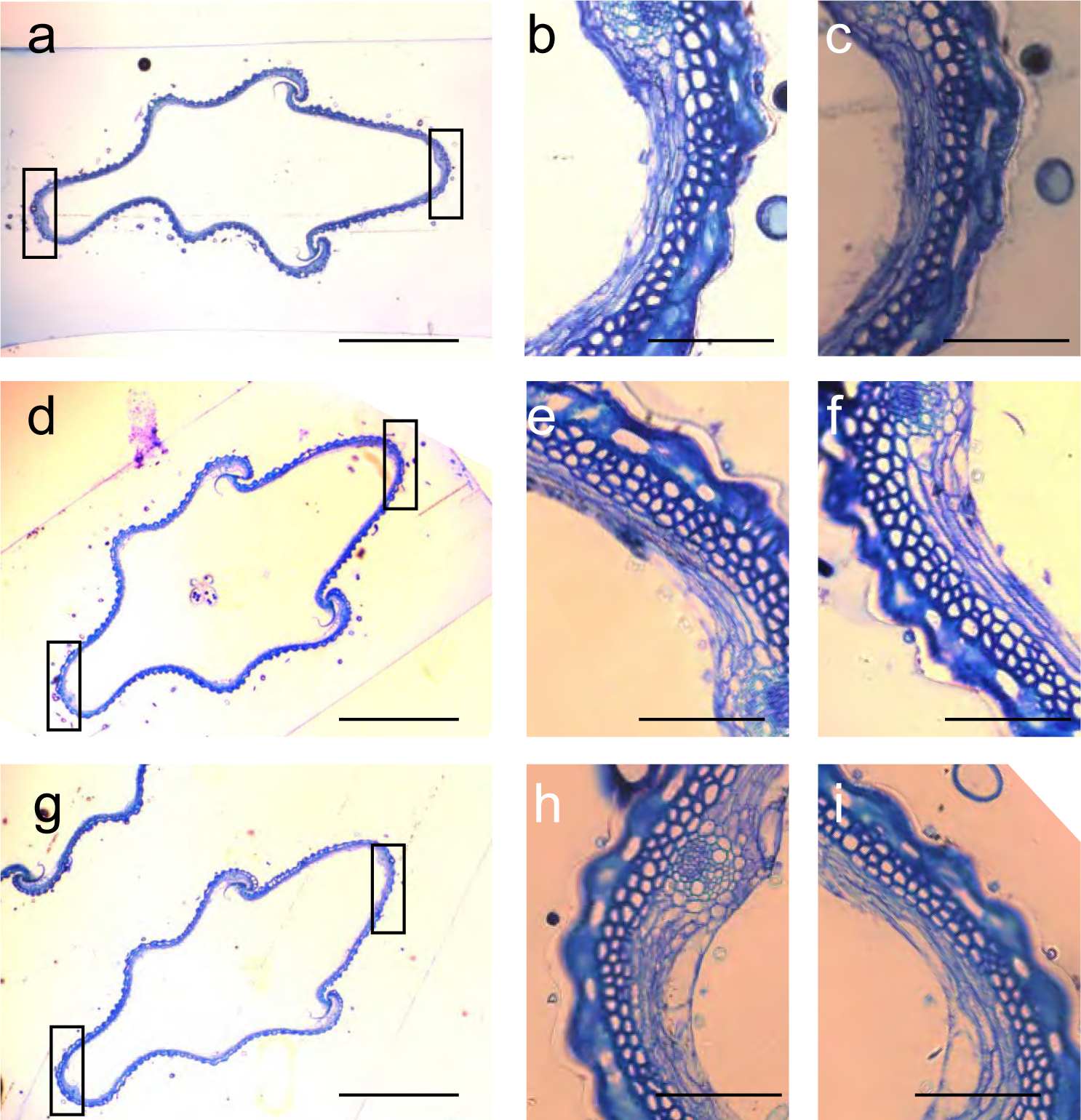
Effect of modulating OsIQD14 expression levels on spikelet hull cells **a-i,** Cross-sections of the spikelet hulls of WT (a-c), *iqd14*-c (d-f) and p35S::*OsIQD14*(g-i) (bar = 1mm). Magnified views of the left column are shown in the middle and right columns (Bar = 0.1mm).

**Supplementary Figure 4.**
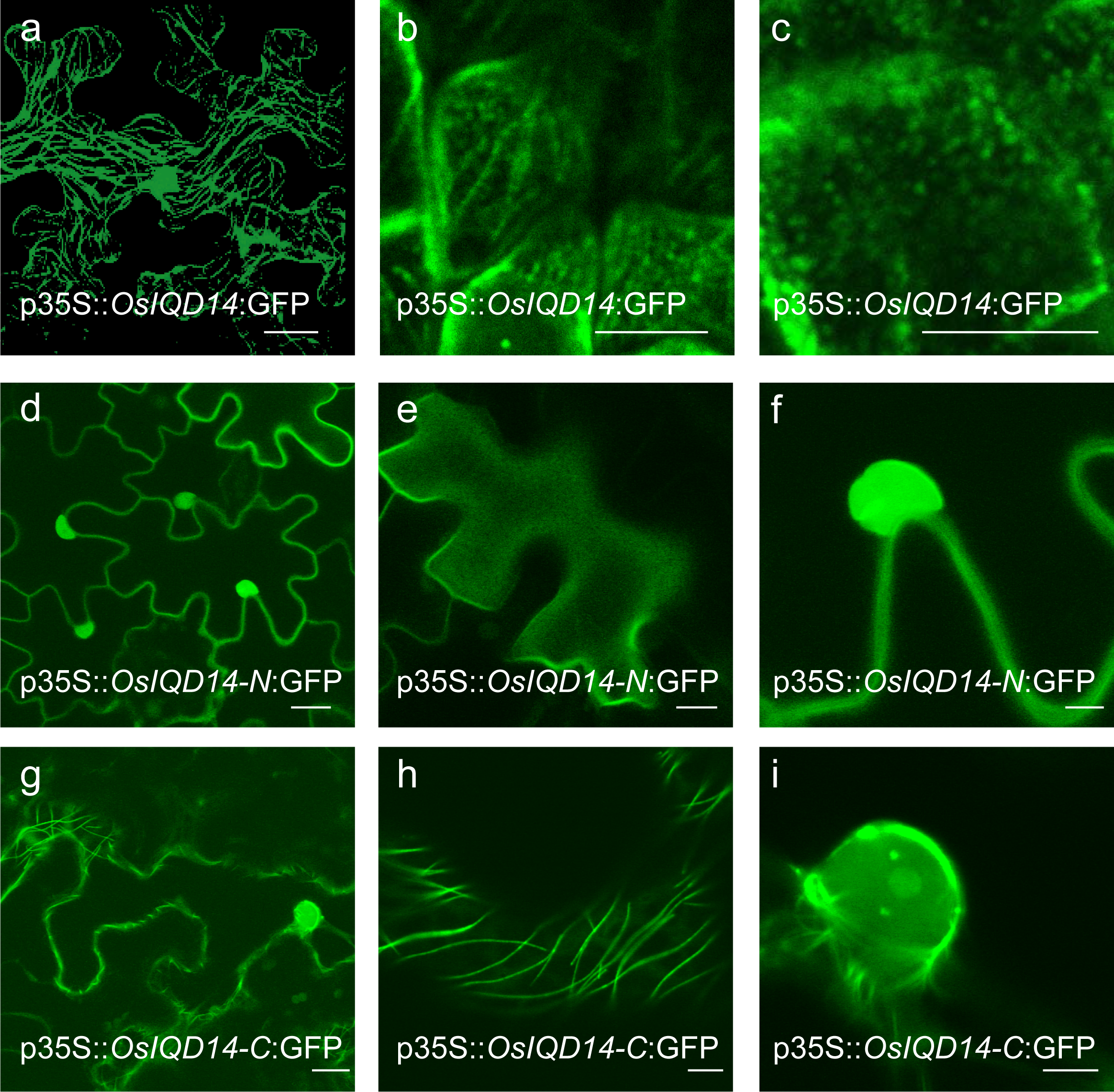
Protein localization of OsIQD14 **a,** Subcellular localization of 35S::*OsIQD14*:GFP in *N. benthamiana* leaf epidermis cells (Z-stack projection; Bar = 20 μm). **b-c,** 35S::*OsIQD14*:GFP is localized to microtubules (left) and depolymerized by oryzalin treatment in *Arabidopsis* root cells (Bar = 5μm). **d-f,** The N-terminal region of OsIQD14 fused to GFP is localized to the nucleus (Bar = 20μm) in *N. benthamiana* epidermis cells. **g-i,** The C-terminal region of OsIQD14 fused to GFP is localized to microtubules (Bar = 20μm). **e, f** and **h, i** show the enlarged zone close to plasma membrane and nucleus (Bar = 10μm or 5μm respectively).

**Supplementary Figure 5.**
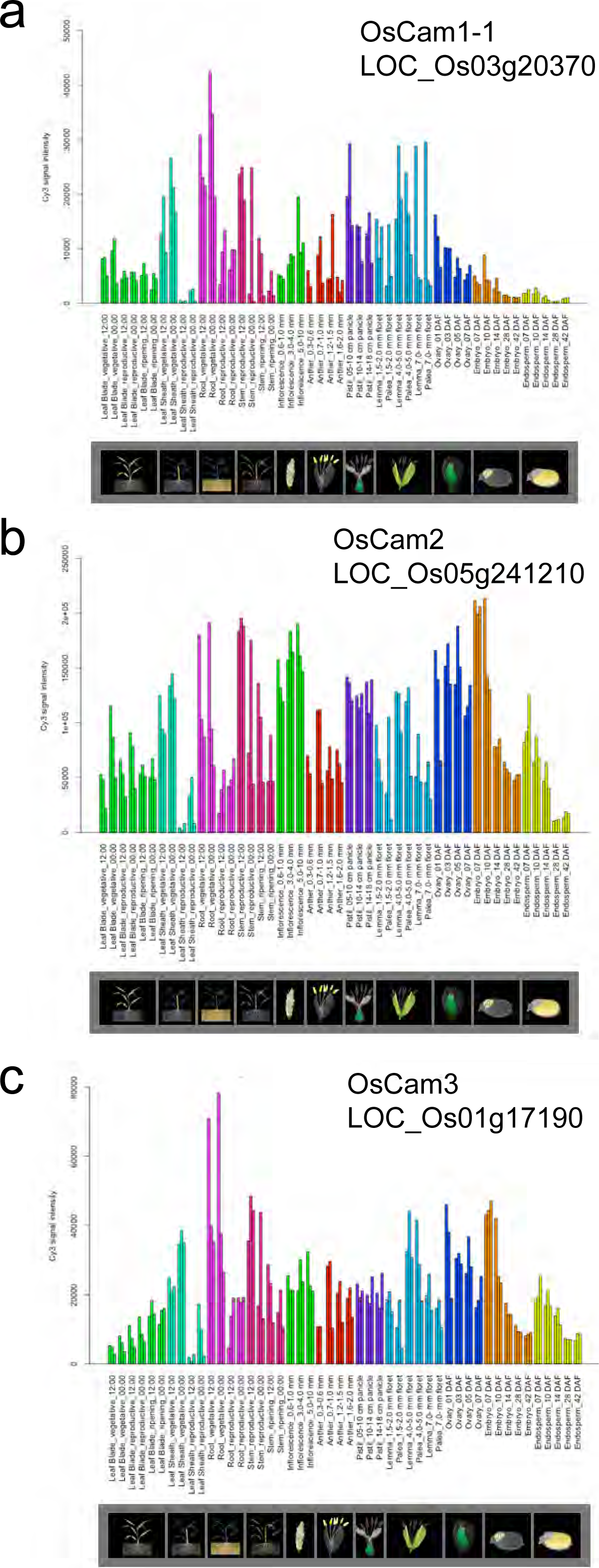
Expression pattern of *OsCaM1,2* and *3* in rice tissue. **a-c,** *OsCaM1,*2 and 3 digital expression levels in leaf, root, inflorescence, anther, pistil, lemma, palea, embryo and endosperm tissues of rice.

**Supplementary Table S1.**
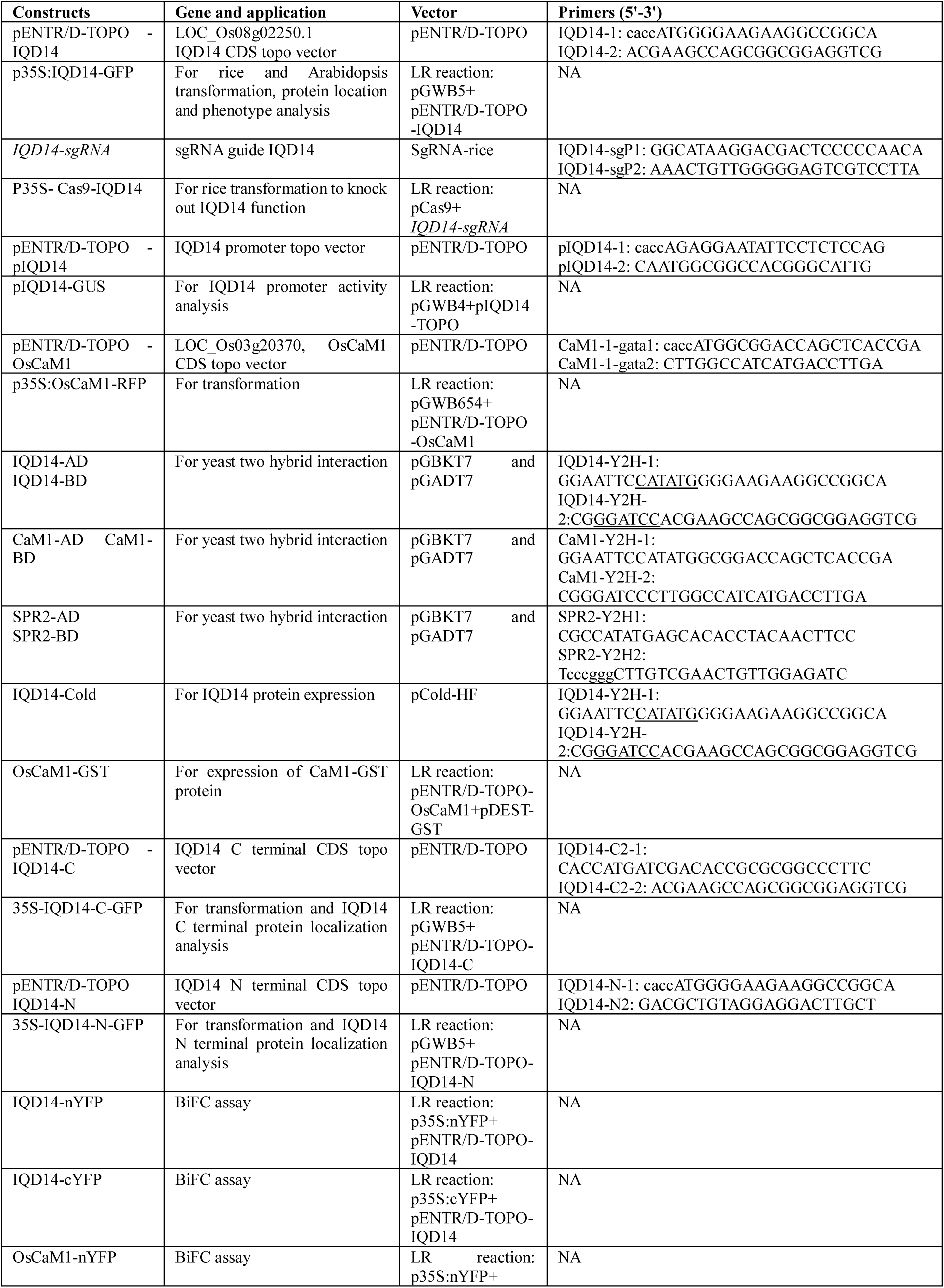
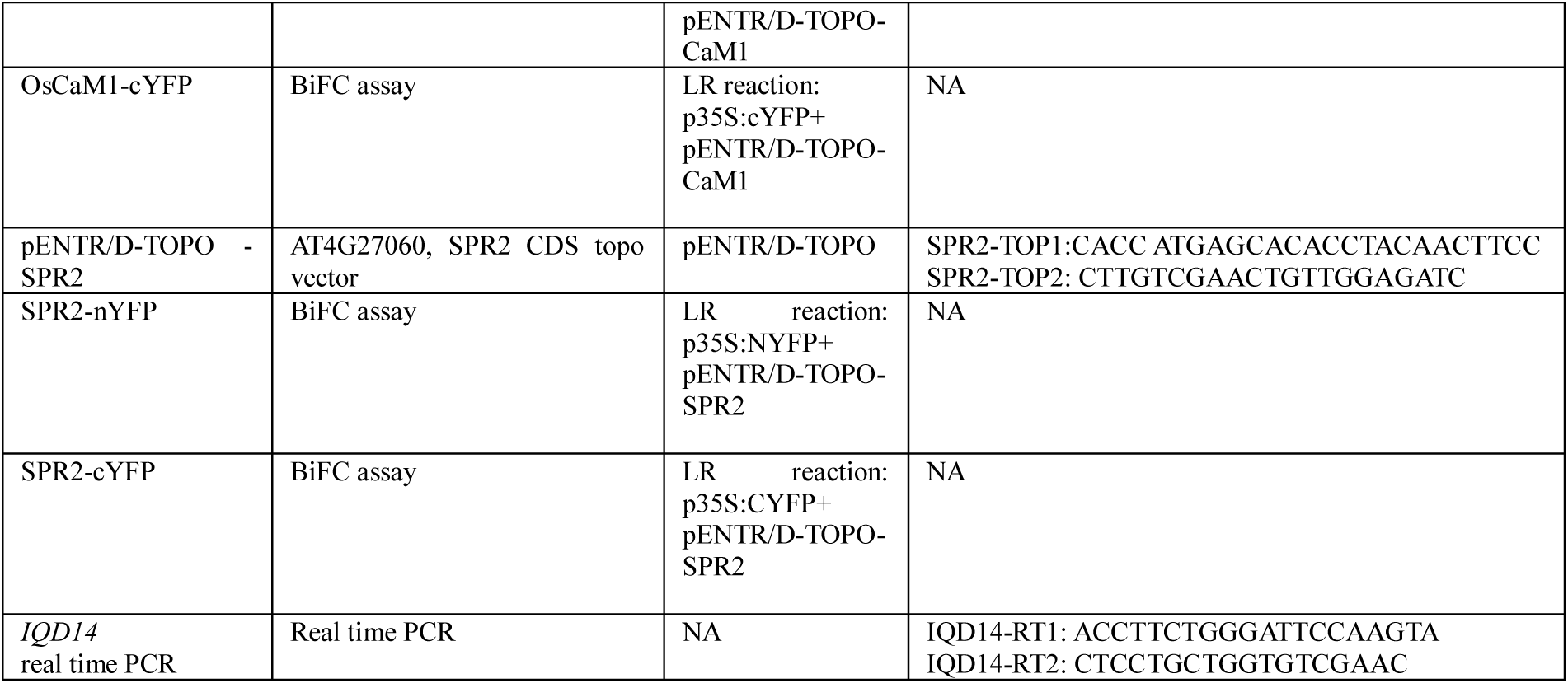
Sequences of the used primers in this study. Added restriction enzymes are underlined.

